# T vector velocity: A new ECG biomarker for identifying drug effects on cardiac ventricular repolarization

**DOI:** 10.1101/418277

**Authors:** Werner Bystricky, Christoph Maier, Gary Gintant, Dennis Bergau, Kent Kamradt, Patrick Welsh, David Carter

## Abstract

We present a new family Tr*X* of ECG biomarkers based on the T vector velocity (TVV) for assessing drug effects on ventricular repolarization. Assuming a link between the TVV and the instantaneous change of the cellular action potentials, drugs accelerating repolarization by blocking inward (depolarizing) ion currents cause a relative increase of the TVV, while drugs delaying repolarization by blocking outward ion currents cause a relative decrease of the TVV.

Evaluating the published data from two FDA funded studies, the Tr*X* effect profiles indicate increasingly delayed electrical activity over the entire repolarization process for drugs solely reducing outward potassium current (dofetilide, moxifloxacin). For drugs eliciting block of the inward sodium or calcium currents (mexiletine, lidocaine), the Tr*X* effect profiles were consistent with accelerated electrical activity in the initial repolarization phase. For multichannel blocking drugs (ranolazine) or drug combinations blocking multiple ion currents (dofetilide + mexiletine, dofetilide + lidocaine), the overall Tr*X* effect profiles indicate a superposition of the individual Tr*X* effect profiles.

The parameter Tr40c allows separating pure potassium channel blocking drugs from multichannel blocking drugs with an area under the ROC curve (AUC) value of 0.90, CI = [0.88 to 0.92]. This is significantly larger than the performance of J-T_peak_c (0.81, CI = [0.78 to 0.84]) using the published data from the second FDA study. Further performance improvement was achieved by combining the ten parameters Tr10c to Tr100c in a logistic regression model, resulting in an AUC value of 0.94.

The TVV based approach substantially improves assessment of drug effects on cardiac repolarization, providing a plausible and improved mechanistic link between drug effects on ionic currents and overall ventricular repolarization reflected in the body surface ECG. TVV may contribute to a better assessment of the proarrhythmic risk of drugs beyond QTc prolongation and JT_peak_c.

## Introduction

Drug effects on ion currents affecting the cardiac ventricular repolarization are well understood on the cellular level. On the ECG level, QTc prolongation is an established surrogate marker for Torsade-de-Pointes (TdP), and was introduced as an electrocardiographic biomarker standard for pro-arrhythmic risk assessment via regulatory pathways in 2005 [1]. Since that time, no drugs have been withdrawn from the market due to unexpected induction of TdP, demonstrating that QTc shows good sensitivity in identifying potentially dangerous compounds. However, a major point of criticism relates to the well-known lack of specificity of QTc, which may result in early discontinuation of promising candidate drugs.

Since pro-arrhythmic drugs can affect ECG morphology, a number of repolarization biomarkers have been proposed for the detection of waveform morphology abnormalities [2, 3] including T-wave duration measures [4], area or amplitude-based measures [5], measures based on a vectorcardiographic representation of the ECG [6, 7, 8], or measures that capture symmetry, flatness or notching of the T wave [9]. However, the predictive value of changes in ECG morphology has not been established [1], and little is known about how such morphological parameters are linked to the electrophysiological process at the cellular level [10].

Recently, members of the Comprehensive *in vitro* Proarrhythmia Assay (CiPA) initiative [11] have completed two FDA-sponsored studies which addressed (1) the effect of ion channel blocks on various ECG biomarkers [12, 13] and (2) differentiation of QTc prolonging drugs with TdP risk (elicited by blocking the outward potassium current) from QTc prolonging drugs with minimal proarrhythmic risk (characterized by a balanced block of inward and outward ion currents) [14, 15]. These studies will be referred to hereafter as (1) Study A and (2) Study B respectively. The latter is of particular practical interest since block of additional ionic currents may compensate for the reduced outward current resulting from hERG/iKr current block to subsequently reduce the extent of QTc prolongation and risk of arrhythmia. The studies demonstrate that a block of the hERG/iKr current prolongs both the early phase of repolarization, as quantified by the J-T_peak_ interval, and the late repolarization phase (T_peak_-T_end_ interval). In contrast, late sodium or calcium inward current blocks preferentially affects the early phase of repolarization, as quantified by shortening only the J-T_peak_ interval. The authors conclude that the J-T_peak_ interval represents the best biomarker currently available with respect to differentiating pure hERG/iKr current block from multi-ion channel blocks [15].

In this paper, we present a new family of vectorcardiographic-based biomarkers that link drug effects observed on surface ECGs to drug effects at the level of ion currents and action potentials. The new biomarkers are based on the T vector velocity (TVV), the speed at which the heart’s dipole vector evolves along its spatial trajectory during ventricular repolarization. We define the parameters Tr*X* as the time that it takes to reach certain percentages *X* of the total trajectory length, and combine the drug-induced changes on the heart rate corrected parameters Tr10, Tr20,.., Tr100 in so-called Tr*X*c effect profiles.

Given that the heart’s dipole vector at a fixed moment in time represents the instantaneous potential gradient generated by the action potentials (APs) of all ventricular myocytes, we suggest that the speed at which the heart dipole vector’s trajectory evolves over time is an integral measure of the changes in the cellular repolarization. At the J point, the majority of cardiac myocytes will be in the plateau phase (phase 2) of repolarization. Any pharmacologic intervention with ion channel blocking drugs provoking a steeper downslope of the plateau will hasten final reconstitution of the resting potential. Hence, such change of the AP can be considered as acceleration of the repolarization process. On the other hand, any drug causing a flatter downslope in AP phases 2 or 3 would constitute a delay of the repolarization process. Such alterations in the steepness of the AP slope would be mirrored in the instantaneous spatial velocity of the T vector trajectory, and captured in the Tr*X*c effect profile. Thus, the effect profile reveals the presence, extent and relative phase of drug-induced alterations in the timing (delays or accelerations) in the ventricular repolarization process. We suggest that, for a given drug and drug concentration, this information constitutes a fingerprint of its effect on repolarization. Tr*X*c effect profiles extend repolarization assessment beyond QTc and the T_peak_ fiducial point to a quasi-continuous view onto the entire repolarization process.

In the remainder of the paper, we present our proposed method and the results of its application to the data of the two FDA-sponsored studies. A performance comparison with the existing biomarkers J-T_peak_ and QTc is given. The findings support our hypothesis, and suggest a promising role for this approach in the rating and ranking of a drug’s pro-arrhythmic potential, and in differentiating multi-channel blocks from pure hERG/iKr current blockades.

## Material and methods

### Study data

We use data from two FDA-sponsored studies that are publicly available from the Physionet website [16]. The first study (Study A) [12, 17] contains 5,232 ECGs of 22 healthy subjects partaking in a randomized, double-blind, 5-period crossover clinical trial. Its aim was to compare the effects of four known QTc prolonging drugs (ranolazine, dofetilide, verapamil, and quinidine, each affecting repolarization via different effects on cardiac ionic currents) versus placebo on electrophysiological and other clinical parameters.

The second study (Study B) [14, 18] contains 4,211 ECGs from a randomized, double-blinded, 5-period crossover clinical trial in 22 healthy subjects. It addresses the electrophysiological responses to hERG/iKR current blocking drugs with and without the addition of blockade of either late sodium or calcium current blocking drugs. The 5 treatment periods include dofetilide alone, mexiletine with and without dofetilide, lidocaine with and without dofetilide, moxifloxacin with and without diltiazem, and placebo.

The studies A and B were approved by the FDA Research Involving Human Subjects Committee and the local institutional review board. All subjects gave written informed consent [12, 14].

In both studies, continuous 12 lead ECGs were recorded at a sampling rate of 500 Hz, from which triplicate 10-second ECGs were extracted at each timepoint using the Antares Software (AMPS LLC). The studies contain J-T_peak_ measurements, corrected for heart rate according to [12]. We used these values for comparison with our TVV-based parameters.

### ECG processing

All published ECG files were analyzed with eECG/ABBIOS (AbbVie, Inc.’s proprietary, validated, ECG analysis system) in a semi-automated manner. ECGs were reviewed to identify artifacts, abnormal heartbeats, and unreliable automated annotations. ECGs of concern were manually reviewed to identify and annotate a minimum of 3 heartbeats per ECG with unaffected T-waves and consistent T annotations. The average RR intervals and the Fridericia corrected QT intervals [19] were extracted and used for further analyses.

For each ECG, the 12 lead signals were adjusted to the isoelectric lines, low-pass filtered (bidirectional Bessel filter with 36 Hz), and exported as an annotated ECG file in HL7 format (aECG) with 500 Hz sampling frequency, including the P, Q, J, and T_end_ annotations. The aECG files were loaded into, and further processed using the R system [20].

20 ECGs of Study A contained isolated leads that were strongly corrupted by noise. Those individual leads were set to zero. Since our approach aims at characterizing spatiotemporal properties of the cardiac repolarization process, we reconstructed the vectorcardiogram (VCG) by means of the inverse Dower transformation [21], and extracted the T-loop by limiting the VCG to the repolarization time interval [J + 20ms; T_end_]. The rationale for choosing the Dower matrix is described in the supporting information section. Only normal sinus beats were considered in the further analysis.

### Calculation of TVV and T vector trajectory quantiles Tr*X*

We estimated the TVV by separately calculating the temporal derivatives 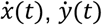 and 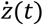 for the three VCG components and combined them as 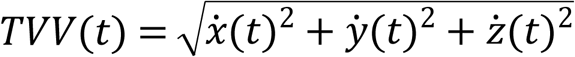. We obtained the temporal derivatives by means of a differentiating Savitzky-Golay polynomial smoothing filter (R method sgolayfilt, package *signal*, version 0.7-6) of order 3 and filter length of 31 samples, equivalent to a time window length of 60 msec.

Integration of the TVV over a time interval corresponds to a cumulative summation of all spatio-temporal changes of the T vector trajectory, and yields the length of the T vector trajectory in the given time span. Normalizing the total T vector trajectory length to 1 allows interpreting the time course of the integrated TVV as a distribution function for the trajectory’s time course. The X% quantile of this distribution corresponds to the time elapsed until X% of the total T vector trajectory length have been reached. To assess drug effects throughout the entire repolarization phase, we used the 10%, 20%, 30%, 40%, 50%, 60%, 70%, 80%, 90%, and 100% trajectory quantiles as parameters, and named them *Tr*10, *Tr*20, …, *Tr*100. For each ECG, we calculated the T vector trajectory quantile *TrX*as the average of the corresponding quantiles of the normal beats.

Since repolarization is known to be strongly affected by heart rate, the trajectory quantile parameters were corrected for heart rate by fitting a linear mixed effects model (R package *lme4*, version 1.1-13) of the form *TrX*∼ 1 + *RR* + (1 + *RR*|*Subject*) to all drug-free data in Studies A and B. The heart rate corrected trajectory quantiles were calculated as *TrXc* = *TrX*– *slope*∗ (*RR*– 1000), where *slope* represents the model’s fixed effect estimate, and RR the ECG’s average beat interval measured in milliseconds.

Fig 1 illustrates derivation of the T vector trajectory quantiles with the ECGs of subject 1017 from Study A under placebo and under quinidine treatment where colors indicate the relative time points. The upper row displays the T vector loops in 3-dimensional space, indicating normal diurnal variation in the placebo column, and quinidine induced T loop morphology changes in addition to circadian effects. Small dots indicate the start of the T vector trajectories, which are 20ms after the J-point. The second row displays the time courses of the T vector strength, which would be the base signals for T_peak_ measurements as applied in the FDA studies A and B (note the flattening and trend toward notching for quinidine). The third row displays the time course of the TVV, and the fourth row the cumulated and normalized TVV over time, which is identical to the relative vector trajectory length at a given time. The horizontal lines denote the 30% and 70% trajectory quantiles. Note that the cumulated TVV curves are nicely aligned with stable relative positions under placebo, but intersect under quinidine. Notably, in the first hours after quinidine dosing corresponding to high quinidine plasma concentrations, the 30% quantiles are decreased, while the 70% quantiles are increased. The time axes in Fig 1 are corrected for heart rate according to Fridericia’s formula.

**Fig 1.**
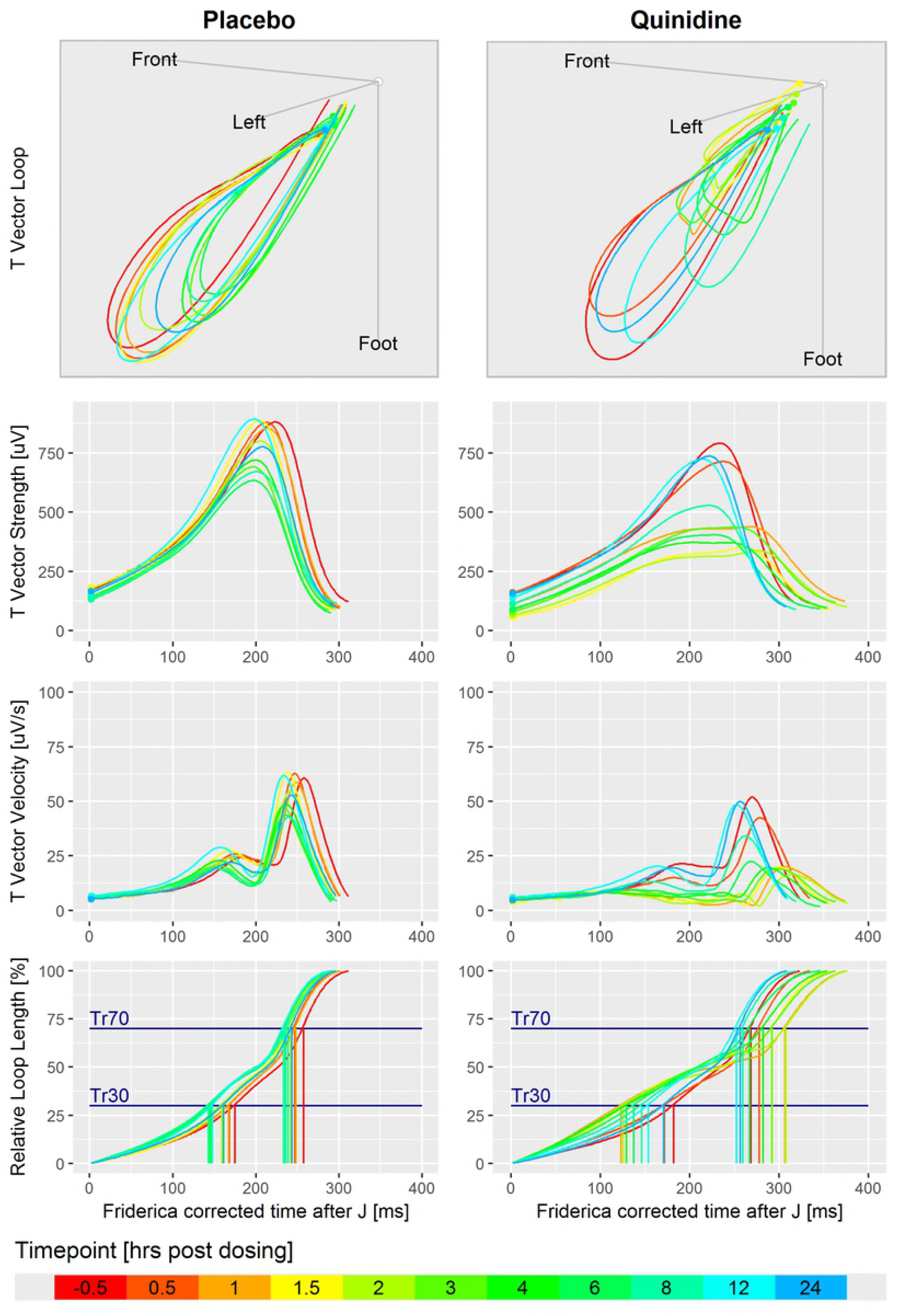
Illustration of the derivation of the TVV and the T vector trajectory quantiles. Displayed are ECGs of subject 1017 in Study A under placebo (left column) and under quinidine treatment (right column). Colors indicate the relative time with respect to dosing. 1^st^ row: T vector loops in the 3-dimensional space. Each loop is the average loop of the normal heartbeats in the first replicate at the given timepoint. The coordinate system is located in the center of the heart with the axes pointing to the body’s front, left side and to the foot. 2^nd^ row: Time courses of the T vector strength. 3^rd^ row: Time courses of the TVV. 4^th^ row: Cumulated and normalized TVV over time, that is the relative loop length reached at a given time, with 30% and 70% trajectory quantiles.

### Evaluation of drug effects

The T vector trajectory quantiles Tr*X*c described above are the basis for our evaluation of drug effects on ventricular repolarization. In order to quantify a drug effect, single-delta parameter values were calculated as Δ*P*(*t,TR*) = *P*(*t,TR*) – *P*(*t*_0_*,TR*), with *P*(*t,TR*) as the average parameter value of the replicate ECGs at time t under treatment TR for a given subject, and t_0_ as the baseline time point.

Double-delta parameter values were calculated as ΔΔ*P*(*t,Drug*) = Δ*P*(*t,Drug*) – Δ*P*(*t,Placebo*).

We modeled the dependencies of the T vector trajectory quantile parameters on the drug concentrations through the mixed effects models ΔΔ*P*∼ 0 + *C* + (0 + *C*│*Subject*) (for single drugs in Study A) respectively ΔΔ*P*∼ 0 + *C*1 + *C*2 + *C*1 ∗ *C*2 + (0 + *C*1 + *C*2│*Subject*) (for multiple drugs in Study B), where ΔΔ*P* is the placebo corrected change from baseline of the parameter P, and C, C1, C2 are the drug concentrations. The zero terms in both models enforce the regression line or plane to intersect with the origin.

The drug effect was calculated as the model’s prediction at representative plasma drug concentrations. For Study A, we chose the following drug concentrations: dofetilide: 2500 pg/ml, quinidine: 1500 ng/mL, ranolazine: 2000 ng/mL, verapamil: 100 ng/mL. For Study B, the representative drug concentrations were determined as the geometric mean drug concentration from the time points where both drugs have been administered (see Table 1). The prediction’s 95% confidence intervals were calculated through a bootstrap simulation.

**Table 1.** Representative plasma drug concentrations in Study B

| Drug combination | Drug | Concentration (ng/mL) |
| --- | --- | --- |
| **Dofetilide + Mexiletine** | Dofetilide | 1.43 |
|  | Mexiletine | 1170 |
| **Dofetilide + Lidocaine** | Dofetilide | 1.28 |
|  | Lidocaine | 1826 |
| **Moxifloxacin + Diltiazem** | Moxifloxacin | 6984 |
|  | Diltiazem | 71.8 |

For assessing the combined effects of mexiletine with dofetilide, we fitted the model to the pooled data from the dofetilide alone treatment period and the period with combined mexiletine plus dofetilide treatment. The same approach was used for assessing the effects of lidocaine combined with dofetilide. In the assessment of the effect of diltiazem combined with moxifloxacin, we omitted the interaction term *C*1 ∗ *C*2 because no data exist for pure diltiazem treatment, and because the regression surfaces for the model including the interaction term revealed implausible distortions. Thus, the predictions of the pure diltiazem effects are model extrapolations and should be interpreted with an appropriate level of caution.

The effects of a drug on the 10 trajectory quantiles Tr10c to Tr100c (together with the corresponding 95% confidence intervals) constitute what we call a drug effect profile (of Tr*X*). A drug effect profile reveals during which phase, and to what extent, a drug delays or accelerates the repolarization process. We derived effect profiles for all drugs in Study A and the various drug combinations in Study B, and examined the effect profile characteristics for their consistency across both studies, and for their plausibility from a physiological perspective.

### Classification of pure hERG/iKR current block versus multichannel block

In order to verify the utility of the T vector trajectory quantile parameters in differentiating pure hERG/iKR current block (group I) from multichannel block (group II), we used the data from Study B following the methods as described in [15]. ECGs from all timepoints at which selective hERG/iKr current blockers had been dosed (dofetilide and moxifloxacin alone arms in Study B) were considered for group I. ECGs from all timepoints with additional administration of late sodium blockers (mexiletine with dofetilide and lidocaine with dofetilide arms in Study B) were assigned to group II.

For each of the 10 trajectory quantiles separately, and for various parameter combinations additionally, we fitted logistic regression classification models (Method glm, R package *stats*, version 3.4.1) to the double-delta parameter data and assessed their performance in separating group I ECGs from group II ECGs by calculating the area under the receiver-operating curve (AUC). The 95% AUC confidence interval was estimated using a stratified bootstrap technique with 2000 replicates (R package *pROC*, version 1.10.0).

### Relation of T vector trajectory quantiles to J-T_peak_ and QT

Since the 100% trajectory quantile Tr100 represents the time interval from J + 20ms to the end of the T wave, we expect that the drug effects as captured by this parameter are closely related to those obtained from the QT interval (resp. QTcF). To verify this, we calculate the drug effects of both parameters for Study A and Study B, and compare them visually.

Keeping in mind that T_peak_ is typically located roughly in the middle of the J-T_end_ interval, we likewise expect a relation between the J-T_peak_ interval and the central T vector trajectory quantiles. This is addressed by visually comparing the drug effects of J-T_peak_ to Tr50 and Tr60 for both studies A and B. In order to be consistent with existing literature, the heart rate corrected J-T_peak_ intervals were taken from the published study data.

## Results

For Study A, Fig 2 displays the drug effects of dofetilide, quinidine, ranolazine, and verapamil in terms of an exposure-response relationship on the (double-delta) 30%, 50%, 70%, and 100% T vector trajectory quantiles. For dofetilide (Fig 2 left column) the slopes in all four graphs are positive. The remaining three compounds show a negative slope for Tr30c (Fig 2, row 1, columns 2-4). Fig 3 displays the effect profiles of all Tr*X*c parameters using representative drug concentrations as described in section 3.4 and indicated as vertical grey lines in Fig 2. Please note that negative effect profile values relate to negative slopes in the exposure-response relation, and positive values correspond to positive slopes. Drug effects on the 10% quantile were negative for all four drugs. Dofetilide displayed a continuously increasing effect profile up to the 100% quantile. The quinidine effect profile was negative up to the 30% quantile with a stronger increase in the middle phase, and slightly reduced growth in the later repolarization phase. The ranolazine effect profile was negative up to the 40% quantile, increased in the mid repolarization phase and stayed close to constant till end of repolarization. Verapamil’s effect profile was slightly negative up to the 40% quantile, and stayed close to zero up to the 100% quantile.

**Fig 2.**
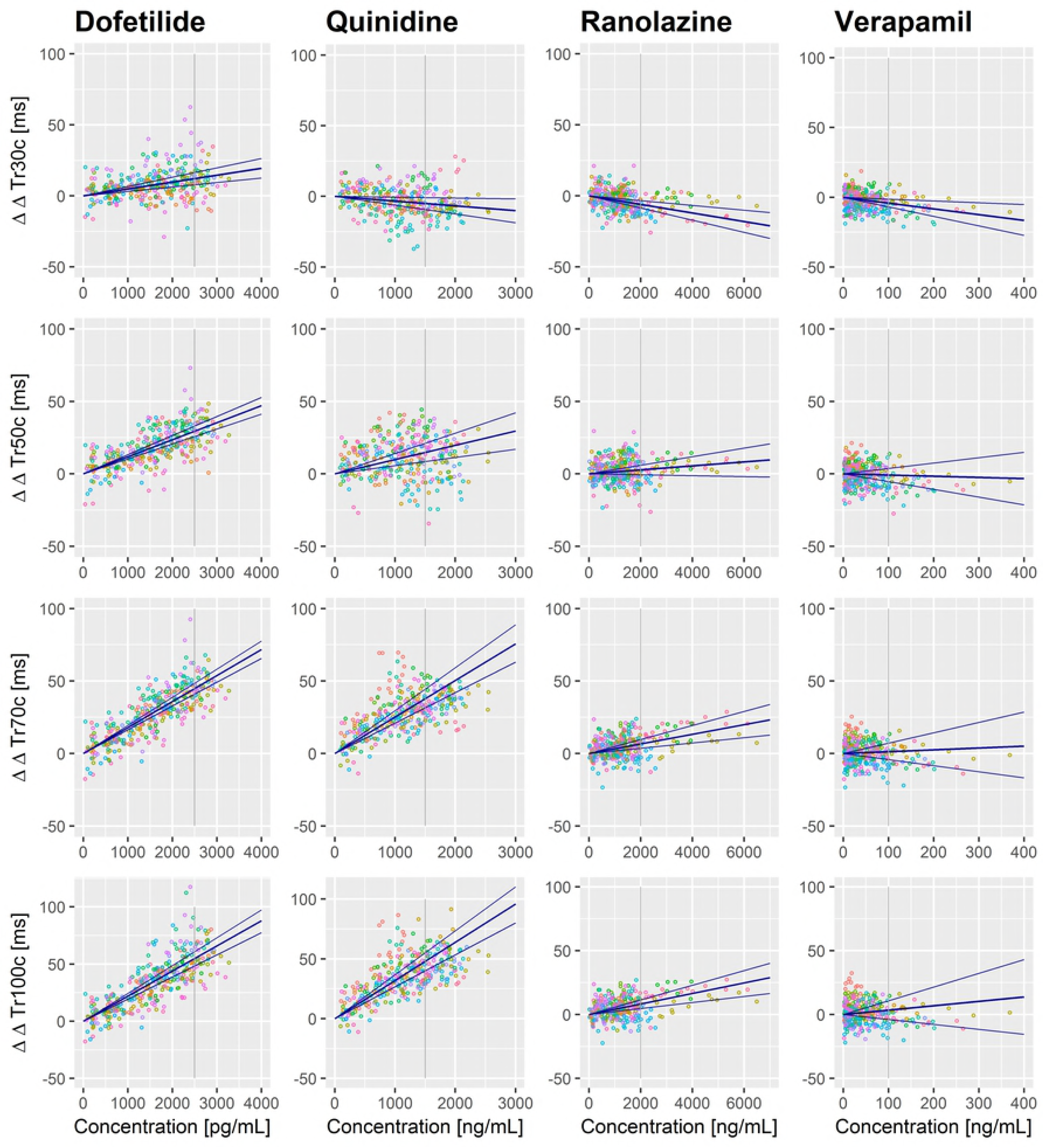
Exposure response in Study A. Effects of dofetilide, quinidine, ranolazine, and verapamil determined as exposure-response relation of the plasma drug concentrations on the double-delta parameters Tr30c, Tr50c, Tr70c, and Tr100c. The blue lines indicate the estimated slope with its 95% confidence interval. The colors of the data points denote individual subjects. The grey vertical lines denote the drug concentrations used for calculating the drug effect profiles in Fig 3.

**Fig 3.**
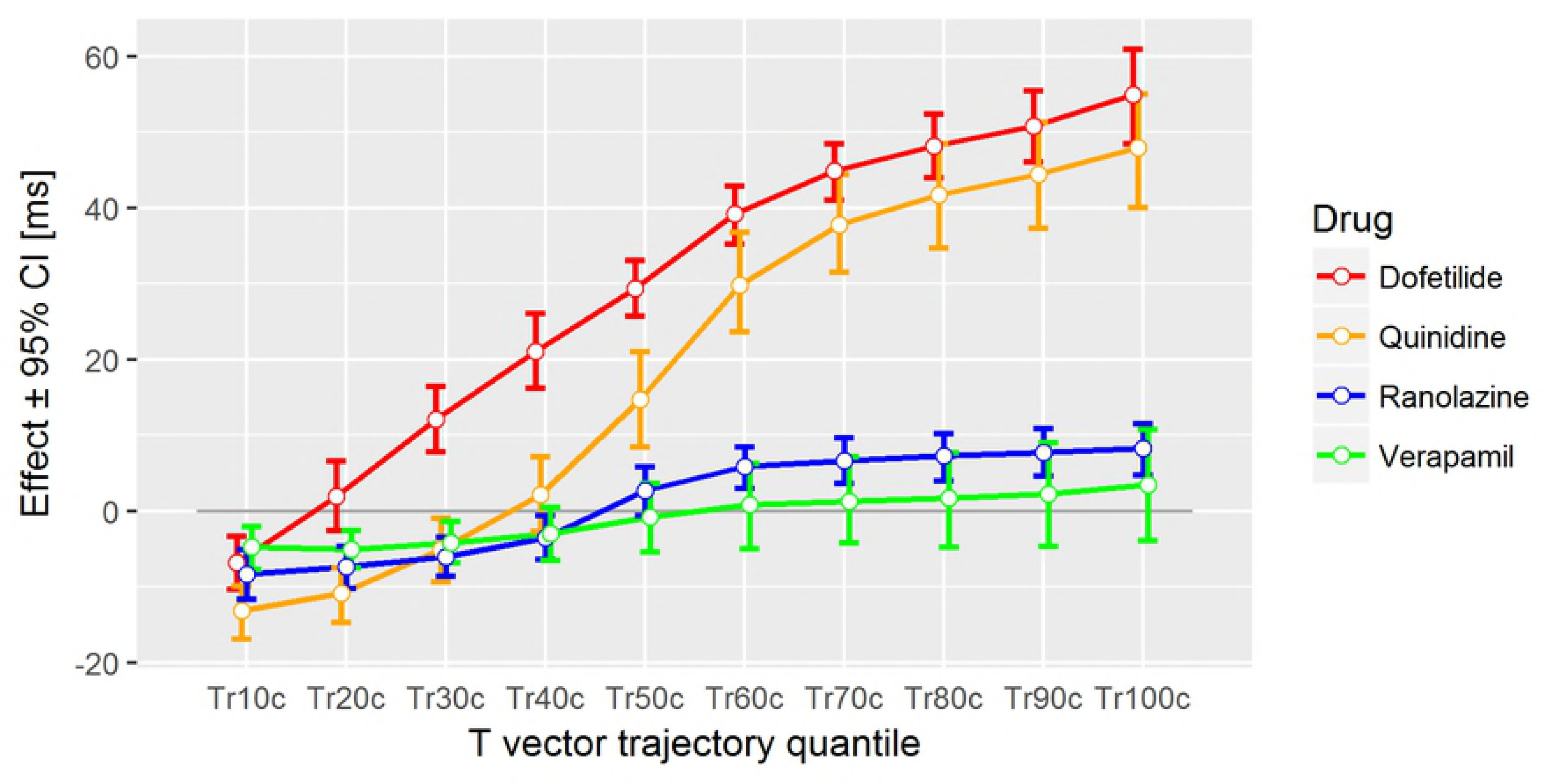
Drug effect profiles in Study A.

The effect on the y-axis denotes the placebo-corrected changes from baseline of the T vector trajectory quantiles in milliseconds (corrected for heart rate) with their 95% confidence intervals at the following drug concentrations: dofetilide: 2500 pg/mL, quinidine: 1500 ng/mL, ranolazine: 2000 ng/mL, verapamil: 100 ng/mL.

For Study B, the effect profiles for the heart rate corrected T vector trajectory quantiles of the various drug combinations are displayed in Fig 4. The drug concentrations used for calculating the drug effect profiles are listed in Table 1. The effect profiles of pure mexiletine and pure lidocaine were very similar in shape, showing a close to zero effect for the 10% quantile, a continuous decreased effect profile up to the 40% quantile, and keeping the negative effect up to the 100% quantile, where mexiletine had a more pronounced negative effect value than lidocaine. The effect profile of pure dofetilide was very similar in shape to the effect profile in Study A. The combinations of mexiletine plus dofetilide and of lidocaine plus dofetilide generated sigmoid-like effect profiles, which were similar in shape to the ranolazine effect profile in Study A: The 10% to 40% quantiles were negative with a transition to positive in the mid-repolarization phase, and a slightly increasing trend up to the 100% quantiles. The effect profile of moxifloxacin alone was completely positive with constantly increasing values from the 10% to the 100% quantile. Combining moxifloxacin with diltiazem slightly increased the late-repolarization related quantiles, and slightly decreased the early-repolarization related quantiles.

**Fig 4.**
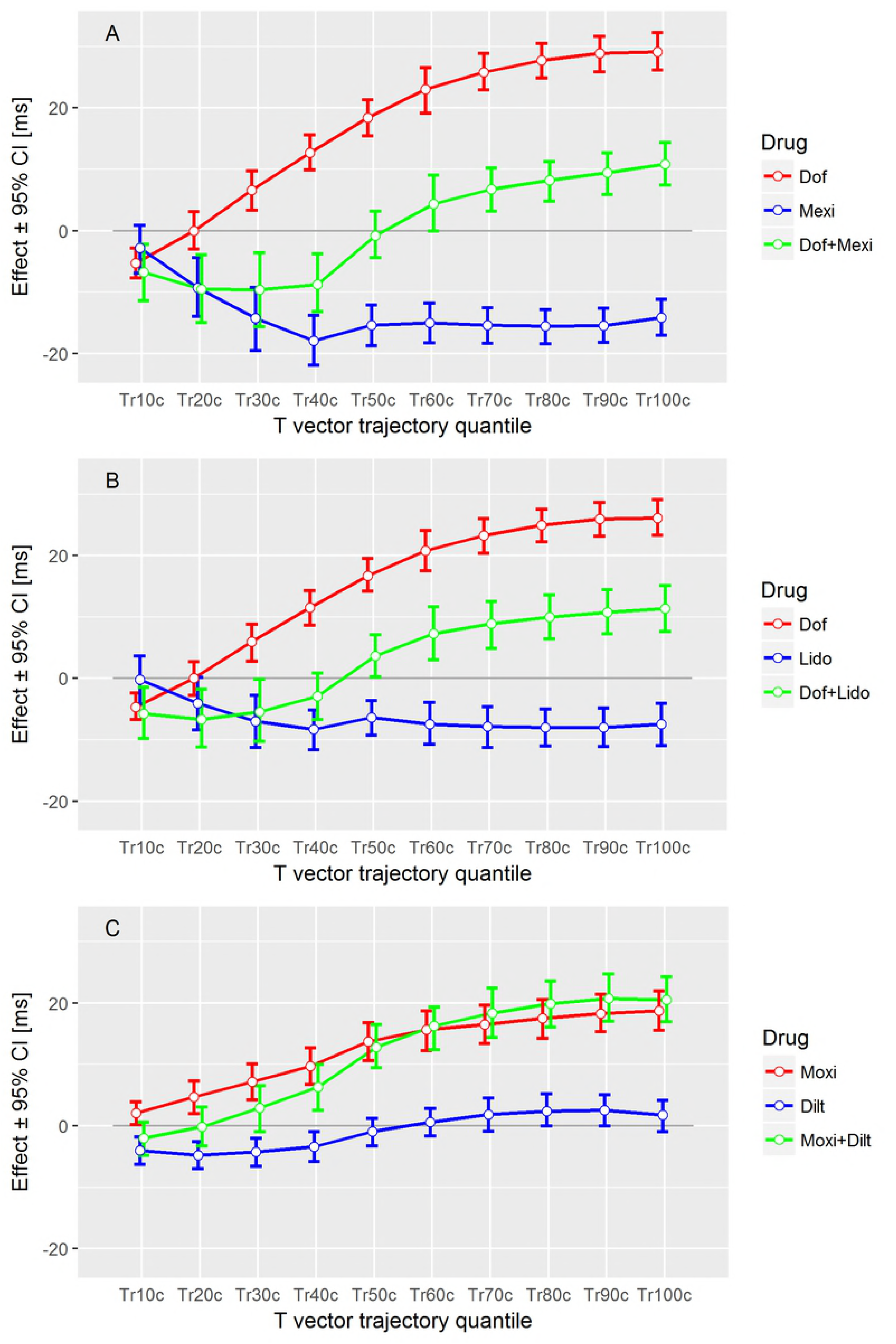
Drug effect profiles in Study B. The effect on the y-axis denotes the placebo-corrected changes from baseline of the T vector trajectory quantiles in milliseconds (corrected for heart rate) with their 95% confidence intervals at the drug concentrations as given in Table 1.

Figs 5 and 6 demonstrate that the double-delta changes of the 100% T vector trajectory quantiles are comparable in size and precision to the double-delta changes of QTcF for all treatments. The highest correlation with the published J-T_peak_ data was observed for the 50% and 60% T vector trajectory quantiles. Table 2 compares the median values and interquartile ranges of the three parameters for the drug-free cases, and shows that the effect size of J-T_peak_ lies between the 50% and 60% quantiles. This also holds for the majority of the cases under treatment (Figs 5 and 6). Notably, most confidence intervals for J-T_peak_c were larger than those of the 50% and 60% T vector trajectory quantiles.

**Table 2.** Distribution of J-T_peak_ compared to T vector trajectory quantiles.

| Parameter | Median (ms) | Interquartile range (ms) |
| --- | --- | --- |
| $J-T_{peak}$ | 221 | [206 to 235] |
| Tr50 | 194 | [180 to 210] |
| Tr60 | 215 | [199 to 231] |

(Note that the T vector trajectory quantiles Tr50 and Tr60 are measured from 20ms after the J point.)

**Fig 5.**
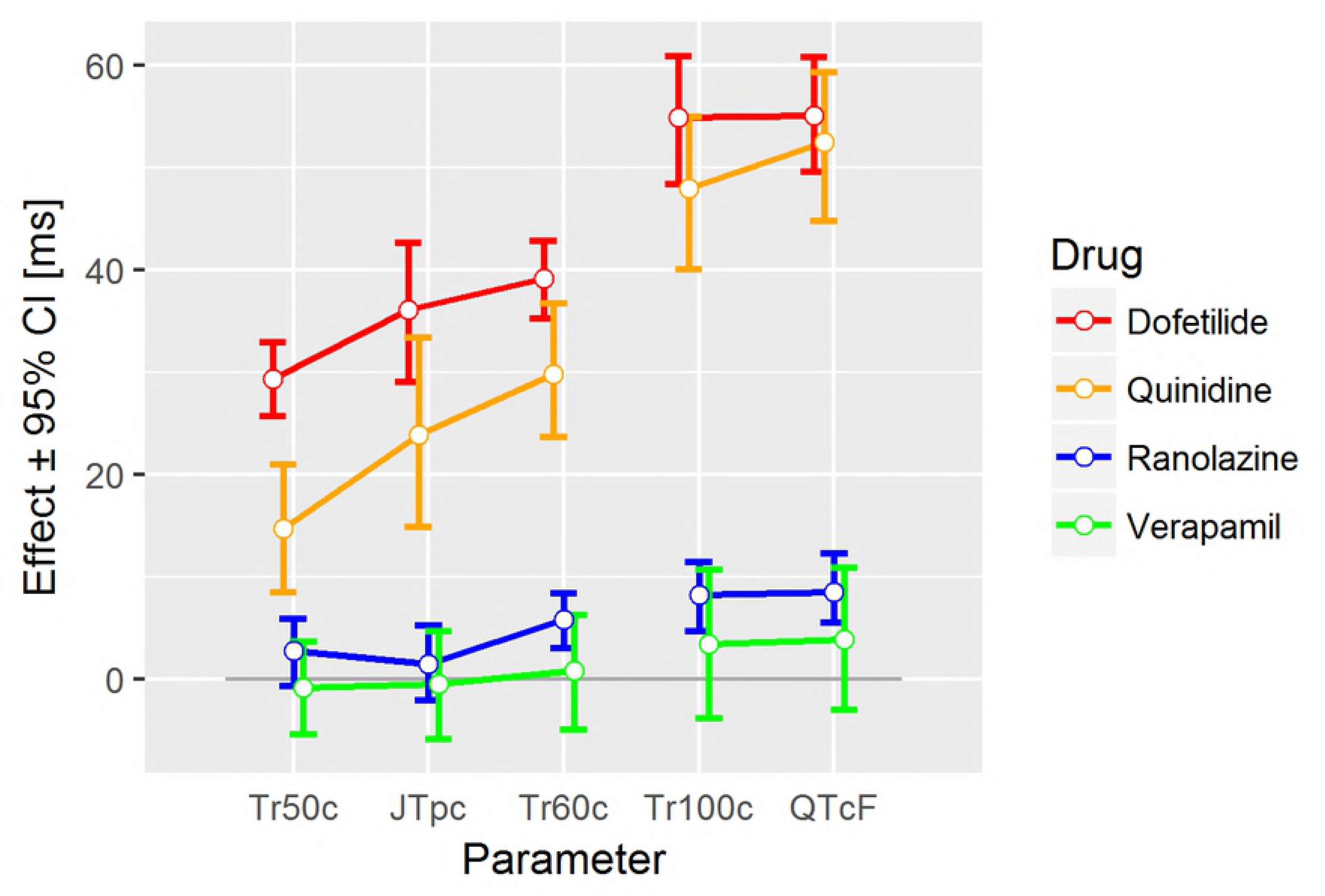
Drug effects on selected parameters in Study A. The drug effects on J-T_peak_c are about between the effects of the 50% and 60% T vector trajectory quantiles (Tr50c and Tr60c), but have larger confidence intervals. The effects on the 100% T vector trajectory quantile (Tr100c) are similar to the effects on QTcF.

**Fig 6.**
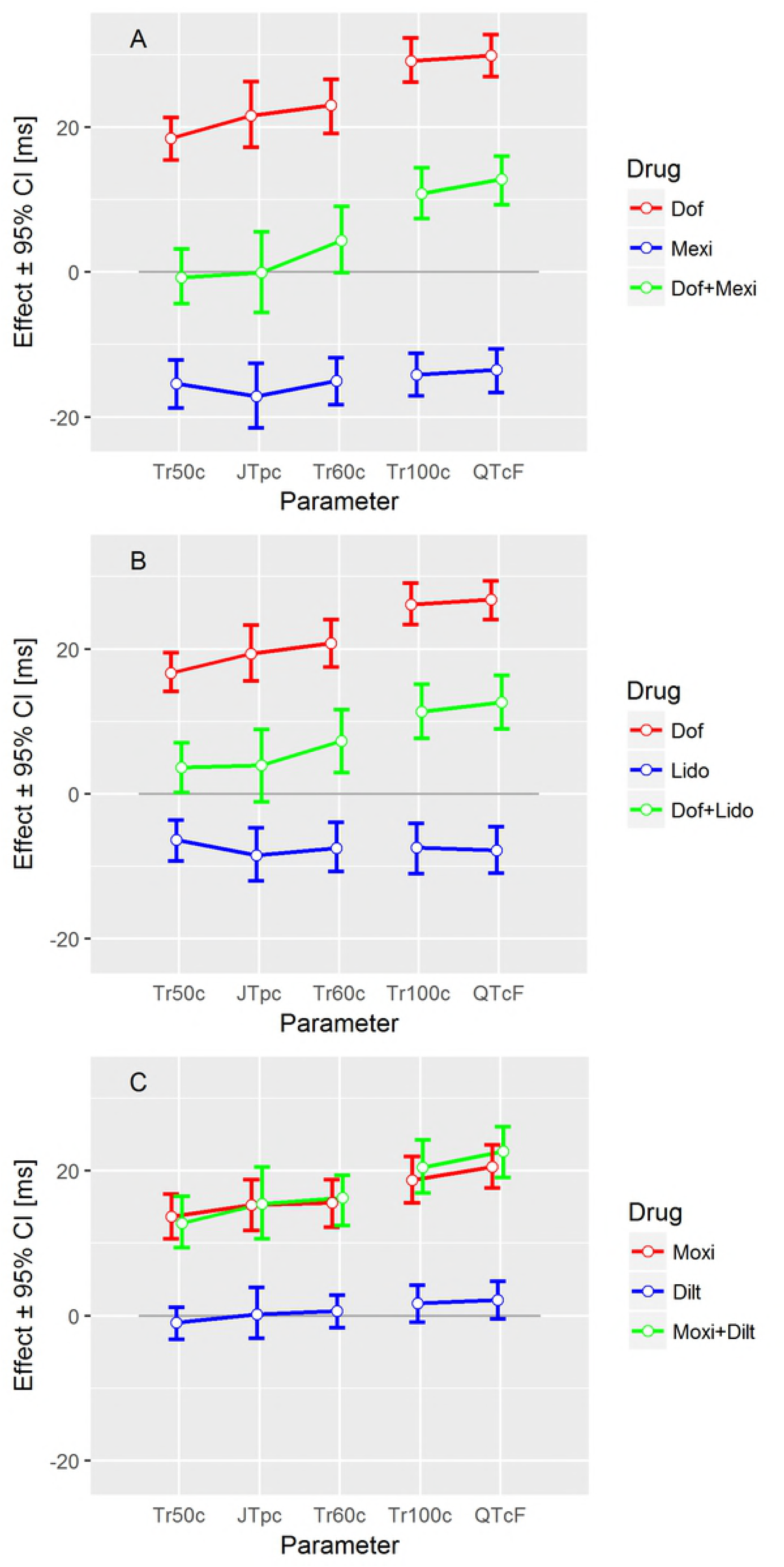
Drug effects on selected parameters in Study B. The drug effects on J-T_peak_c are between the effects of the 50% and 60% T vector trajectory quantiles (Tr50c and Tr60c), but have larger confidence intervals. The effects on the 100% T vector trajectory quantile (Tr100c) are similar to the effects on QTcF.

The AUC values for separating selective hERG/iKr current block versus multichannel block with late sodium current inhibition (Study B) are displayed in Figs 7 and 8. The largest AUC value for a single parameter was observed for the 40% T vector trajectory quantile with 0.90, CI = [0.88 to 0.92] (Fig 7). Combining all 10% step quantile parameters increased the AUC value to 0.94, CI = [0.92 to 0.96] (Fig 8). The AUC value for J-T_peak_c (published data) was 0.81, CI = [0.78 to 0.84], and the AUC value for QTcF using the eECG/ABBIOS generated annotations was 0.73, CI = [0.69 to 0.77].

**Fig 7.**
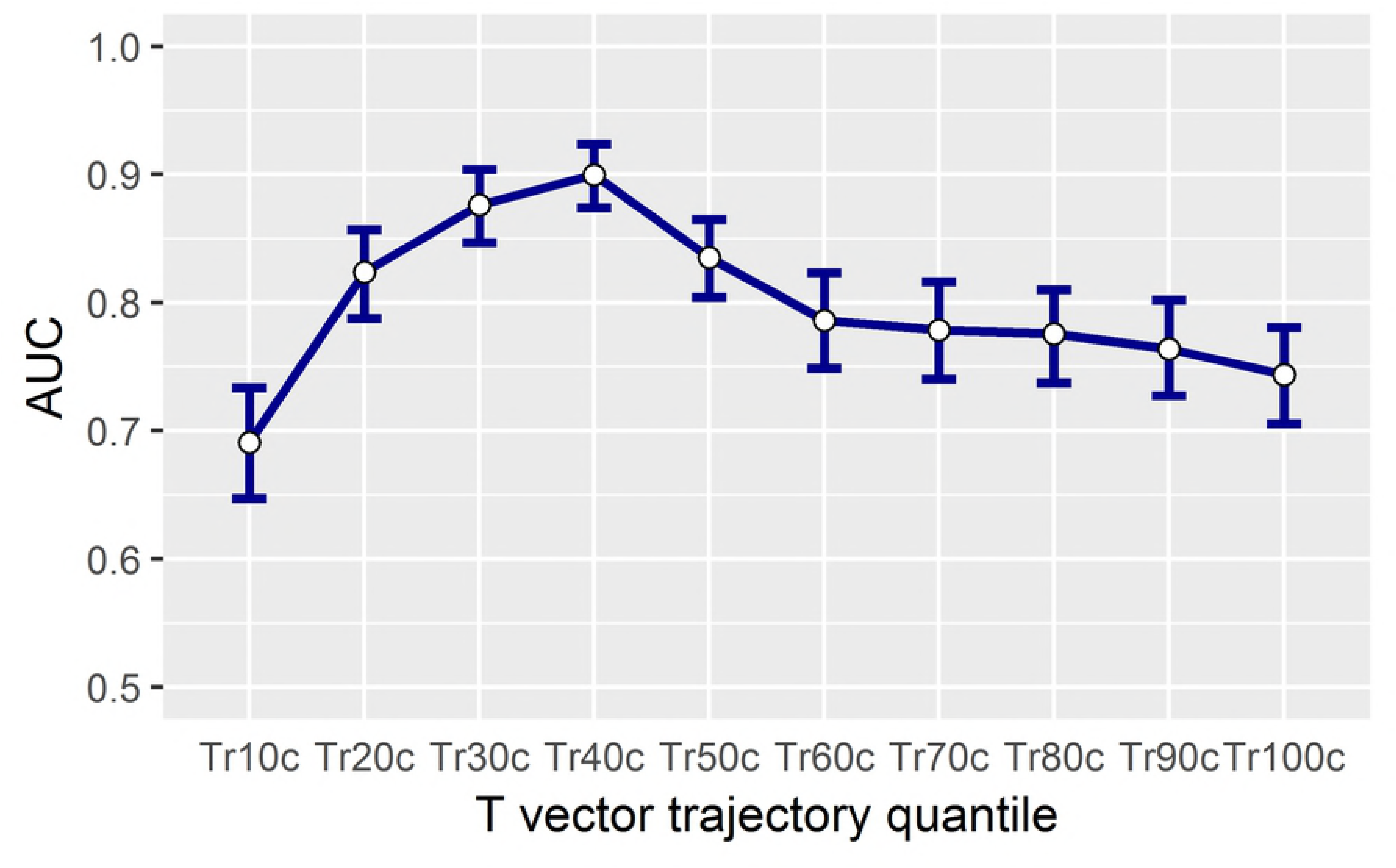
Separation performance of pure hERG/iKr current block versus multichannel block based on individual T vector trajectory quantiles. Separation performance between group I and group II (see section 3.5) is measured by the area under the receiver-operating curve (AUC) with its 95% confidence intervals. The T vector trajectory quantiles Tr10c to Tr100c are corrected for heart rate. Best separation is observed for the 40% quantile Tr40c.

**Fig 8.**
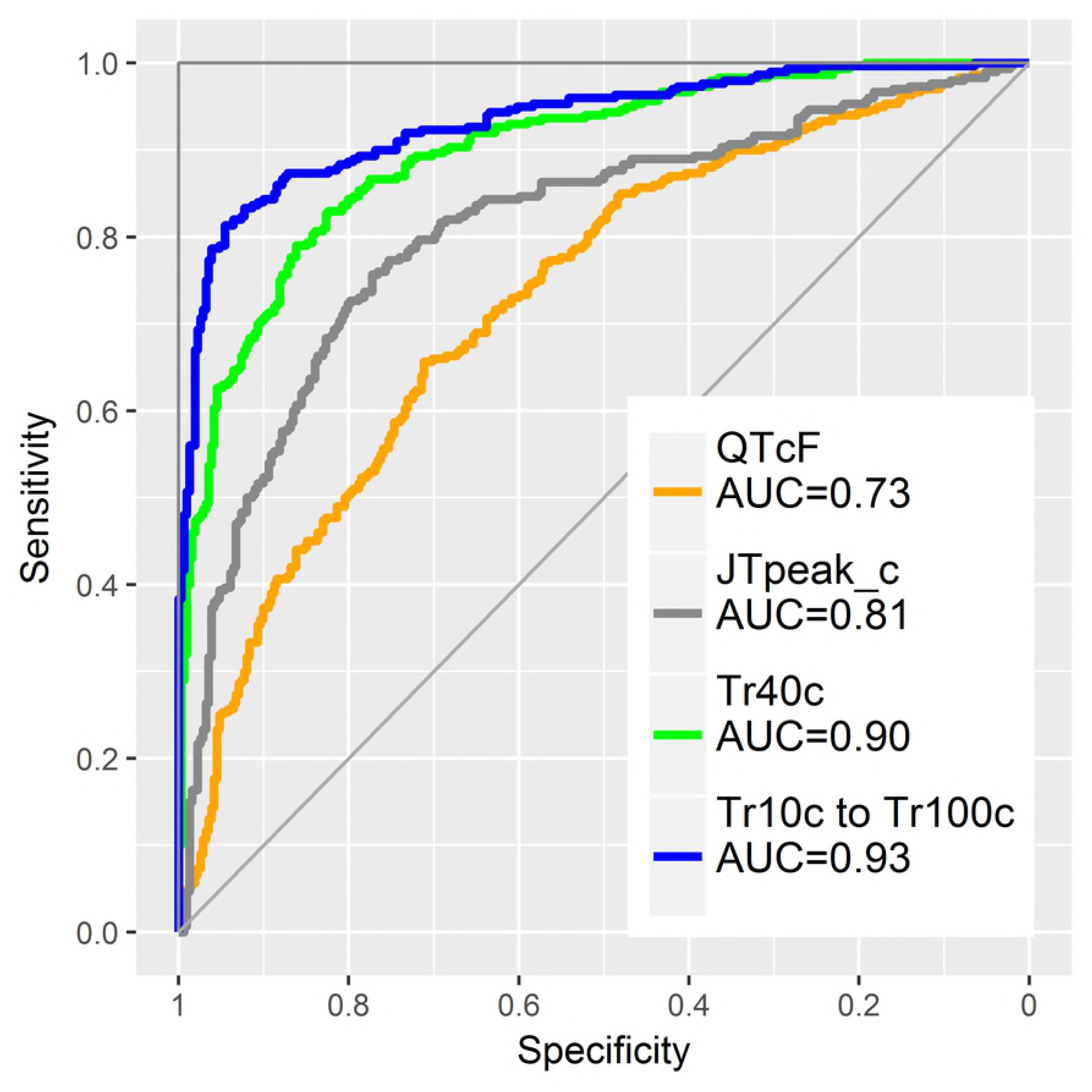
Classification of pure hERG/iKr current block versus multichannel block in logistic regression models. The separation performance (AUC) of the heart rate corrected T vector trajectory quantile Tr40c (green curve) and the combined 10 Tr*X* quantiles (blue curve) are compared to QTcF (orange curve) and to published [15] J-T_peak_c data (grey curve).

## Discussion

### Interpretation of T vector trajectory quantiles

In this study, we used heart rate corrected quantiles of the heart’s dipole vector trajectory along the T wave loop as a new set of ECG biomarkers for assessing drug effects on the repolarization process. The *X*% T vector trajectory quantile Tr*X* denotes the time when *X*% of the total T vector trajectory length has been reached. Arriving earlier at a particular point on the T vector trajectory (as compared to placebo) means that the repolarization process is accelerated in the corresponding phase of repolarization. Hence, the dose-response curve of this placebo-corrected change from baseline quantile shows a negative slope, and its effect profile value is negative. In contrast, an increase in the time required to reach a particular point on the T vector trajectory indicates a delay of the repolarization process, and is associated with a positive slope in the dose-response curve and a positive value in the effect profile. The magnitude of an effect profile value reflects the effect size, i.e. the extent of acceleration or delay, with respect to a given drug concentration.

We propose a link between cellular repolarization changes resulting from blockade of single and multiple ionic currents and the observed Tr*X* effect profiles as illustrated in Fig 9: Inward sodium and calcium ion currents are predominantly active during earlier repolarization, and both contribute to the maintenance of the action potential’s plateau phase. In our simulation based on the O’Hara-Rudy model [22], blocking the late sodium ion channels causes a steeper descent of the AP in phase 2 (Fig 9 emerald curve vs. red curve). This represents an acceleration of the repolarization process and earlier restitution of the resting potential. Phase 3 of the AP appears only marginally affected by pure sodium ion channel block (Fig 9) as the red and emerald curve run almost in parallel in phase 3, i.e. their downslopes have comparable magnitude.

**Fig 9.**
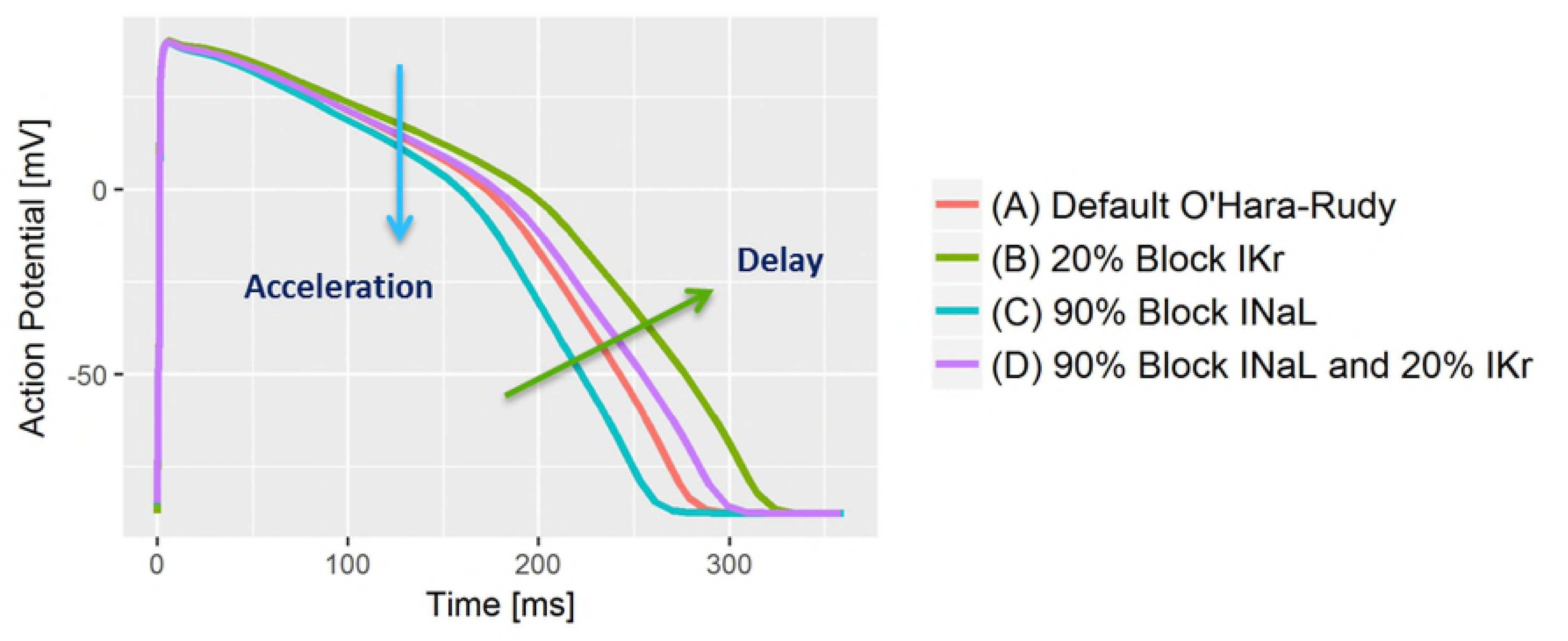
Simulation of blocking of the late Sodium (INaL) and the hERG/IKr currents in the O’Hara-Rudy model. Blocking INaL accelerates the repolarization process by increasing the downslope in AP phase 2. Blocking IKr decreases the downslope of the AP in both, phase 2 and phase 3, resulting in delayed repolarization. In simultaneous blocking, the effects can compensate in phase 2 but a delay in phase 3 is still evident..

We expect that faster intracellular action potential changes are reflected in higher speed of progression along the T vector trajectory. If sodium-block related acceleration happens in phase 2 of the AP, it should increase the TVV in the initial part of the T vector trajectory. Hence, the corresponding lower T vector trajectory quantiles will be reached earlier, resulting in negative effect profile values for the lower quantiles. This phenomenon is observable in all effect profiles of the late sodium current blockers mexiletine (Fig 4A blue line) and lidocaine (Fig 4B blue line), where the lower quantiles Tr10c to Tr40c are continuously decreased. The accumulated level of time lead is maintained over the second part of repolarization with Tr50c to Tr100c remaining at a negative effect profile level comparable to that of Tr40c. This is in accordance with the limited effect of pure sodium ion channel block onto phase 3 of the AP in our simulation (Fig 9). Overall, this indicates that the slight QTcF shortening effect of these two drugs is mainly caused by an acceleration of early repolarization, which then is largely maintained.

Fig 9 also shows that pure hERG/iKr current block causes a reduction of the downslope in the AP in both phase 2 and phase 3 (green curve vs. red curve), corresponding to a delay which affects all parts of the repolarization process. The effect profiles of both pure hERG/iKr current blockers dofetilide (red curves in Figs 3, 4A and 4B) and moxifloxacin (red curve in Fig 4C), consistently reflect this. They show a continuously increasing delay, indicating that the QT prolongation caused by these drugs are effective over the entire course of repolarization.

A combined block of sodium and hERG/iKr currents results in a superposition of the single effects in the AP waveform (Fig. 9 violet curve). The reduced downslope of pure hERG/iKr current block (green curve) is largely compensated for by the additional sodium channel block in phase 2 (Fig 9 violet curve vs. red curve), but is still evident in phase 3 (note the reduced downslope of the violet curve versus the red one). The effect profiles of the corresponding drug combinations dofetilide + mexiletine (Fig 4A, green curve) and dofetilide + lidocaine (Fig 4B, green curve) are well in line with this. Their sigmoid shape indicates a slight, but largely balanced, acceleration of the repolarization process in the early T vector trajectory quantiles Tr10c to Tr40c (associated with the effect of the sodium ion channel block in AP phase 2 and the counteracting hERG/iKr current block), which then becomes increasingly delayed by the preponderance of the hERG/iKr current block effect in AP phase 3.

In the case of mexiletine, the combined effect profile (Fig 4A, green curve) is an approximately linear combination of the pure drug effect profiles (Fig 4A, red curve and blue curve). For lidocaine, the effect profile of the drug combination (Fig 4B, green curve) is slightly more depressed compared to the pure lidocaine effect alone (Fig 4B, blue curve). This is owing to a significant contribution of the interaction term *C*1 ∗ *C*2 in the mixed effects model to all T vector trajectory quantile parameters.

The presented interpretation is further supported by the effect profiles of drugs, which simultaneously affect multiple ion channels. Ranolazine blocks both the hERG/iKr and the late sodium currents, and shows the same sigmoid-shaped effect profile (Fig 3 blue curve) observed for the combinations of the individual channel blockers (Figs 4A and 4B green curves).

Verapamil predominantly blocks the L-type calcium ion channel, mainly affecting AP phase 2, i.e. early repolarization, and to a lesser extent the hERG/iKr current. In line with this, its effect profile (Fig 3, green curve) indicates a slight but significant acceleration of the heart’s electrical activity in the early repolarization, which is compensated for during late repolarization but does not result in significant prolongation of the overall repolarization duration, measured by the QTcF interval.

The effect profile of quinidine (Fig 3, orange curve) as a strong hERG/iKr current blocking drug as well as calcium and sodium current blocking agent indicates significant acceleration in the early repolarization phase and a strong delay in the mid and late repolarization phases. On the other hand, the effect profile of dofetilide (Fig 3, red curve) as a pure hERG/iKr current blocker indicates (with exception of Tr10c) continuously increasing delay throughout repolarization.

As reported in [14], the combination of diltiazem with moxifloxacin in the evening did unexpectedly slightly increase QTcF, despite a slightly reduced moxifloxacin plasma concentration compared to the afternoon (pure moxifloxacin). Our analysis shows that diltiazem slightly accelerates repolarization in the early phase (Fig 4C, green curve for Tr10c to Tr40c), which may be due to its calcium ion channel block. However, diltiazem delays the later repolarization phase, finally increasing the repolarization delay induced by moxifloxacin. Note that the pure diltiazem effect profile (Fig 4C, blue curve) was calculated as an extrapolation of the mixed effects model, and should be interpreted with care.

In summary, extending the lessons learned from the analysis of the J-T_peak_ interval, our results support that block of different ionic currents which are active at different times during repolarization does not have a uniform effect on repolarization characterized using TVV. Blocking of inward ion currents that maintain the plateau phase 2 of the cardiac action potential (late sodium and calcium) causes overall acceleration of the cellular repolarization process, mainly during earlier repolarization. In contrast, blocking of the hERG/iKr current increasingly delays repolarization over its entire course. We propose that systematic analysis of the TVV quantiles permits disentangling the temporal relation and quantifying the relative effect size of ion current blockade at the cellular level from the surface ECG. We suggest that our Tr*X* effect profile concept constitutes an electrocardiographic fingerprint of a drug’s net impact on repolarization, which may indicate effects on different stages of repolarization and different ion channel blockade. To the best of our knowledge, no other surface ECG analysis approach with this promise has been published before.

### Differentiation of pure hERG/iKr current block versus multichannel block

Our results confirm that T vector trajectory quantiles enhance differentiation of multichannel block from pure hERG/iKr current block. In our data, pure hERG/iKr current block consistently manifests as an increasing delay of repolarization with a slightly higher slope between Tr10c to Tr60c compared to Tr60c to Tr100c (Fig 3 and Fig 4, red lines). The presence of additional effective ion channel blocks is revealed by deviations from this profile. According to the previous discussion, the effect of late sodium ion channel block should be particularly evident during phase 2 of the AP. The finding that the “early” T vector trajectory quantiles Tr20c, Tr30c and Tr40c demonstrate the best separation (Fig. 7), and that Tr40c is most effective among all T vector trajectory quantiles to identify additional late sodium block (Fig. 7) is well in line with this assumption. Note that the performance of the 40% quantile of the T vector trajectoryTr40c, reaching an AUC of 0.9 (Fig. 8), is significantly higher than that provided by J-T_peak_ (0.81) representing the best previously published ECG biomarker for that purpose [14, 15]. The potential of the effect profile in identifying multi-ion channel effects is highlighted by a further increase in performance observed when the entire profile is used in the classification process (Fig. 8, AUC 0.94). Well aware of the danger of overfitting a 10-parameter-model to a data set of limited size, we take this improvement only as an additional hint that requires prospective validation.

### Relation of Tr*X* to QT and J-T_peak_

Since QTcF and Tr100c just differ in the starting point (Q versus J+20ms) and in a slightly different correction formula for heart rate, it is not surprising that their effects are largely comparable in magnitude and precision. The slight differences may arise from drug effects on QRS duration, or from systematic differences of our algorithms’ robustness with respect to identification of the J point and the onset of Q. Neither difference appears to be significant in our study.

ECG effects described by the Tr*X* parameters are in line with effects described by parameters, which are derived from the T_peak_ fiducial point: Pure hERG/iKr current blocking drugs delay repolarization through the entire repolarization process, indicated by continuously increasing parameters Tr10 to Tr100. Drugs or drug combinations that additionally block late sodium or calcium ion currents shorten the parameters Tr10 to Tr40. Accordingly, pure hERG/iKr current blocking drugs prolong both the J-T_peak_ and the T_peak_-T_end_ intervals, while J-T_peak_ tends to be shortened by multichannel blocking drugs [23]. However, declaring T_peak_ as cutting point between early and late repolarization is questionable [24], and there is no generally accepted physiologic reason for choosing the T_peak_ to subdivide the repolarization interval for assessment of pro-arrhythmic effects [25]. It has recently been challenged whether a lack of J-T_peak_ prolongation can reliably differentiate between safe and proarrhythmic QT_end_ prolongation [26].

Using a simplified model of VCG generation, we would expect T_peak_ to represent the instant in time when the largest spatial gradient of intracellular potentials exists, i.e. a moment of maximum heterogeneity of the action potentials in all heart cells. From our new parameters, the Tr50c and Tr60c quantiles correspond most closely with J-T_peak_c. For the majority of drugs, the effect size of J-T_peak_c is located between that of Tr50c and Tr60c (Figs. 5 and 6, lines on left side).

The clear trend for narrower confidence intervals of Tr50c and Tr60c, compared to J-T_peak_c (Figs. 5 and 6, lines on left hand side), suggests that the determination of the T vector trajectory quantiles is more robust than that of the T_peak_ position. This is not surprising in view of the demonstrable changes of T wave morphology induced by drugs, including severe flattening or notching. In such a dynamics, small alterations of the intracellular potential distribution may cause significant dislocation of the T_peak_ position, increasing J-T_peak_ parameter variability. We attribute the robustness of the T vector trajectory quantiles Tr*X* to their unique identifiability even in the presence of significant changes in T wave / T loop morphology and consider this an advantage compared to J-T_peak_.

Another conceptual difference between J-T_peak_ and the Tr*X* parameters is that the J-T_peak_ determination is purely based on the T vector’s magnitude, while the Tr*X* parameters capture changes of both the magnitude and the direction of the T vector progression. Since directional changes of the T vector trajectory reflect changes of the distribution of electrical activities in the whole heart, this information may contribute to the performance improvement in differentiating multi-channel blockade.

## Conclusion

The T vector trajectory quantiles and the related effect profiles extend the discrete ECG characteristics T_peak_ and QT to a quasi-continuous view on the repolarization process. They allow detailed description of drug effects on the various ionic currents, and allow better differentiation of single versus multichannel block than J-T_peak_.

The TVV based approach described here enhances characterization of drug effects on cardiac myocyte ion channels, and may help in better assessing the proarrhythmic risk of drugs. The proposed functional linkage between drug effects on cellular levels and the TVV measured on the ECG may help in aligning the information retrieved in the four compounds of the Comprehensive in vitro Proarrhythmia Assay (CiPA).

The new TVV-based parameters will have to be verified on a larger set of subjects and drugs. We hope that a third FDA-sponsored study, which is currently under evaluation (ClinicalTrials.gov, NCT03070470) can be used for this purpose.

## Acknowledgments

AbbVie contributed to the interpretation of data, writing, reviewing, and approval of this publication. WB and CM are contractors at AbbVie.

The authors deeply acknowledge the FDA for conducting the Study A and B, and for making the data available to the scientific community.

